# Interactions between ecological, evolutionary, and environmental processes unveil complex dynamics of island biodiversity

**DOI:** 10.1101/099978

**Authors:** Juliano Sarmento Cabral, Kerstin Wiegand, Holger Kreft

## Abstract

**Aims:** Understanding how biodiversity emerges and varies in space and time is central to ecology and biogeography. Multiple processes affect biodiversity at different scales and organizational levels, hence progress in understanding biodiversity dynamics requires the integration of these underlying processes. Here we present BioGEEM (BioGeographical Eco-Evolutionary Model), a spatially-explicit, process-based model that integrates all processes hypothesized to be relevant for biodiversity dynamics and that can be used to evaluate their relative roles.

**Location:** Hypothetical oceanic islands

**Methods:** The model is stochastic, grid-based, and integrates ecological (metabolic constraints, demography, dispersal, and competition), evolutionary (mutation and speciation), and environmental (geo-climatic dynamics) processes. Plants on oceanic islands served as model system. We used the full model to test hypotheses about emergent patterns at different spatio-temporal scales and organizational levels (populations, species, communities, and assemblages), switching off processes to assess the importance 1) of competition for realistic population and range dynamics; 2) metabolic constraints for endemism and community composition; 3) environmental dynamics and 4) speciation for biogeographical patterns.

**Results:** The full model generated multiple patterns matching empirical and theoretical expectations. For example, populations were largest on young, species-poor islands. Species, particularly endemics, were better able to fill their potential range on small, species-poor islands. Richness gradients peaked at mid-elevations. The proportion of endemics was highest on old, large, and isolated environments within the islands. Species and trait richness showed unimodal temporal trends. Switching off selected processes affected these patterns, and we found most of our hypotheses supported.

**Main conclusions:** Integrating ecological, evolutionary, and environmental processes is essential to simultaneously generate realistic spatio-temporal dynamics at population, species, community, and assemblage level. Finally, large-scale biodiversity dynamics emerged directly from biological processes which make this mechanistic model a valuable ‘virtual long-term field station’ to study the linkages between biogeography and ecology.

## INTRODUCTION

Ecologists and biogeographers have a long-standing interest in explaining how species are distributed in space and time, but disentangling the relative role of various potential mechanisms for the generation and maintenance of biodiversity remains a challenge (Pennisi, 2005). Considering the complex interlinkage between eco-evolutionary processes and environmental dynamics indicated to influence biodiversity, it seems crucial to account for these processes simultaneously (Urban *et al.*, 2016; Cabral *et al.*, 2016). This has been achieved by complex mechanistic models that simulate macro eco-evolutionary processes (e.g. colonization, speciation, extinction) directly at the species and/or community ecological levels to generate the biogeographical patterns of interest (e.g. Gotelli *et al.*, 2009; Colwell & Rangel, 2010). Other models can simulate processes at lower ecological levels, such as propagule dispersal, individual survival, or population establishment, to generate colonization and extinction as emergent processes (e.g. Harfoot *et al.*, 2014; Singer *et al.*, 2016; Urban *et al.*, 2016). This would provide insights across ecological levels and contribute to integration of biogeographical and ecological theories (Evans *et al.*, 2013; Rosindell & Harmon, 2013; Cabral *et al.*, 2016).

The major limitation of biogeographical models that simulate processes at low ecological levels is high model complexity. Integrating multiple processes increases the number of parameters and complicates interpretability (Dormann *et al.*, 2012). These issues can be avoided by investigating multiple emergent patterns at different scales, i.e. pattern-oriented modelling (Grimm & Railsback, 2012). Pattern-oriented modelling allows distinguishing different parameter combinations that generate similar patterns at a given scale by evaluating patterns at other scales. To counter-balance model complexity, a usefully complex model should thus be able to predict patterns at multiple spatio-temporal scales and at different levels of ecological organization (e.g. populations, species, communities). Likewise, a useful study system should be as simple as possible, but still informative across scales and ecological levels. Oceanic islands are suitable study systems because they are small and isolated, have distinct boundaries, occur in large numbers worldwide, and exhibit striking examples of evolutionary diversification (Losos & Ricklefs, 2010; Warren *et al.*, 2015). Accordingly, island research has contributed essential information about eco-evolutionary processes that shape biodiversity (e.g. Ricklefs & Bermingham, 2004; Whittaker & Fernández-Palacios, 2007).

Different processes have been postulated to be important drivers for insular biodiversity. The seminal equilibrium theory of island biogeography (ETIB) emphasizes the roles of area affecting extinction and of isolation affecting colonization (MacArthur & Wilson, 1963), and the importance of these factors has received ample support by many macroecological studies (e.g. Kreft *et al.*, 2008; Triantis *et al.*, 2012; Weigelt & Kreft, 2013). Climate and environmental heterogeneity also exert strong influence on insular biodiversity (Kreft *et al.*, 2008; Hortal *et al.*, 2009; Cabral *et al.*, 2014). Recently, the importance of geo-climatic processes and time has been highlighted (Whittaker & Fernández-Palacios, 2007; Losos & Ricklefs, 2010; Lomolino *et al.*, 2010; Weigelt *et al.*, 2016). In particular, species richness and endemism depend on island age, geological ontogeny, and Pleistocene sea-level changes (Whittaker *et al.*, 2008; Borregaard *et al.*, 2016, Weigelt *et al.*, 2016). Speciation is influenced by speciation mode as well as island area, isolation, and environmental heterogeneity (Whittaker *et al.*, 2008; Kisel & Barraclough, 2010; Rosindell & Phillimore, 2011). These factors and associated processes are summarized by the General Dynamic Model of Island Biogeography (GDM), which assumes a humped trajectory of area, elevation, and environmental heterogeneity over a simplified life span and ontogeny typical of many oceanic islands (Whittaker *et al.*, 2008). The GDM predicts humped temporal trends of species richness, endemic richness, colonization, extinction, and speciation rates (predictions recently updated by Borregaard *et al.*, 2016). Therefore, a mechanistic model focusing on insular biodiversity dynamics should generate those patterns alongside patterns at lower ecological levels, such as species-abundance distributions (Ulrich *et al.*, 2010) and species-area relationships (e.g. Triantis *et al.*, 2012).

This study aims to investigate the relative roles of ecological, evolutionary, and environmental processes for biodiversity dynamics using a process-based BioGeographical Eco-Evolutionary Model (BioGEEM). Our main hypothesis is that the integration of these three main process types is needed to accurately model biodiversity patterns. Oceanic islands served as our model systems. In our model, we extended niche models with low-level ecological processes (e.g. dispersal, population dynamics, competition, and metabolic constraints) and added evolutionary (mutation and speciation) and environmental (changes in area and environmental heterogeneity) processes. We first used the full model to evaluate emergent patterns against theoretical predictions and empirical patterns. We then assessed the relative importance of the simulated processes by switching off submodels (environmental dynamics, speciation, competition, metabolic constraints) and re-evaluating emergent patterns. We hypothesized that 1) competition is necessary to mediate population and range dynamics of competing species via competitive exclusion (Cabral & Kreft, 2012); 2) metabolic constraints are necessary to generate realistic patterns of endemism and community composition along environmental gradients and via life-history trade-offs (Brown *et al.*, 2004); 3) environmental dynamics and 4) speciation are necessary to generate biogeographical patterns as predicted by island biogeography theory (Whittaker *et al.*, 2008; Borregaard *et al.*, 2016).

## MATERIALS AND METHODS

### General model description

We extended a spatially-explicit multi-species model for range dynamics of plants (Cabral & Kreft, 2012) with relevant evolutionary and environmental processes. We implemented environmental dynamics that reflect the growth and erosion phases of oceanic islands (Fig. 1b; Whittaker & Fernández-Palacios, 2007; Whittaker *et al*., 2008). Ecological and evolutionary processes were controlled by metabolic constraints in a hierarchical structure (Fig. 1a). Specifically, processes were linked to environmental variables (local temperature) and species properties (body mass, Brown *et al.*, 2004). Metabolic constraints generated spatial (via local temperature) and interspecific (via body mass) variation in eco-evolutionary processes and accounted for life-history trade-offs, and thus precluded super-organisms (e.g. minimal resource requirement and maximum reproduction). We focused on terrestrial seed plants, but these metabolic constraints are ubiquitous (Brown *et al.*, 2004). Below, we summarize the model. A detailed description following the Overview, Design concepts, and Details protocol (ODD, Grimm *et al.*, 2010) is provided in Appendix S1 in Supporting Information.

**Figure 1.**
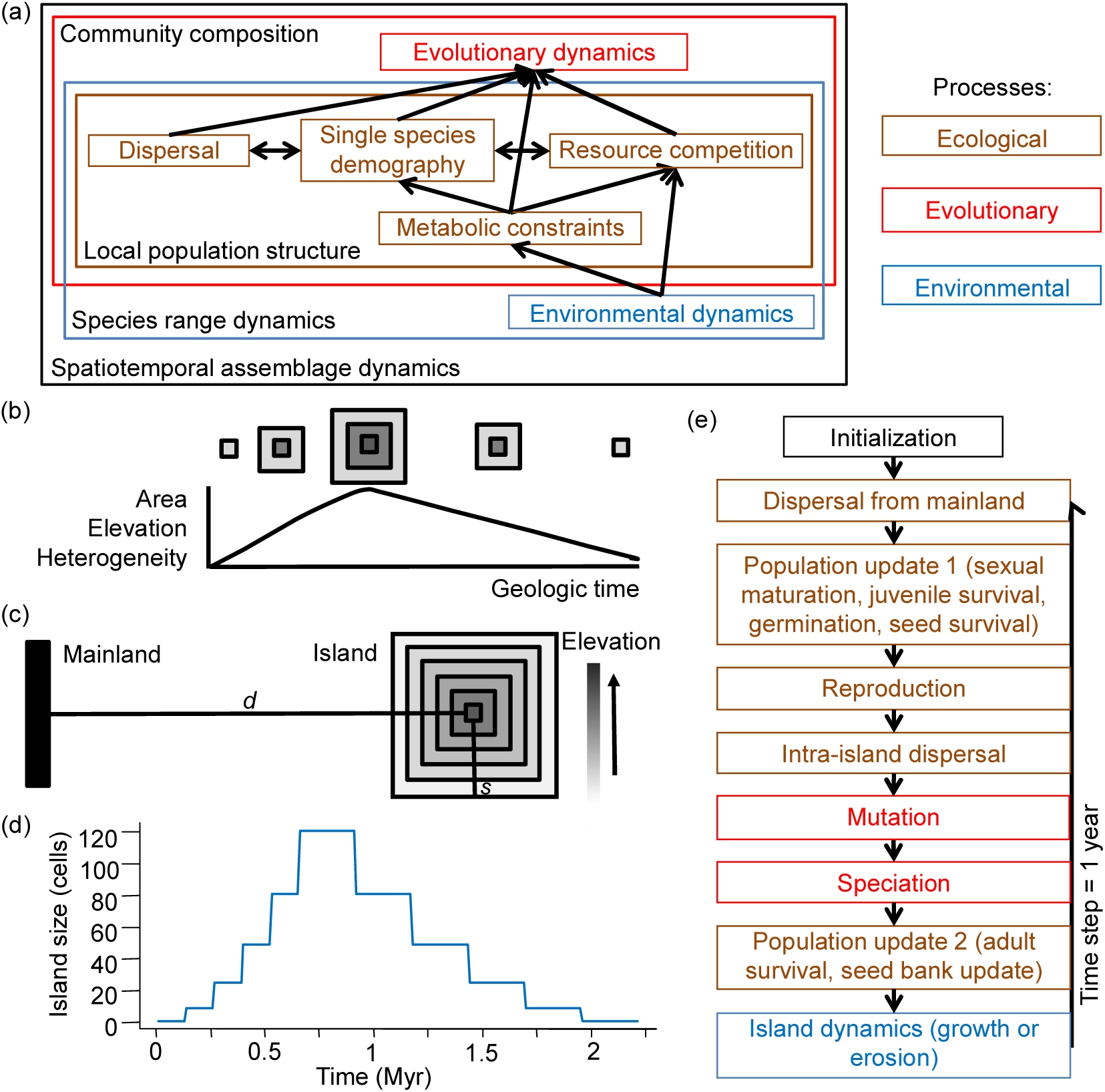
Model framework. (a) Hierarchical structure of simulated processes and emergent patterns. (b) Growth and erosion of hypothetical volcanic islands over geological time. Elevation and environmental heterogeneity are expected to correlate positively with island size and thus to be a humped function of island age. (c) Simulation grid at maximum island size, where *s* is the maximum distance between island centre and edge (*s* = 5 cells). Grid dimensions are described by *s* and distance *d* from the island centre to the mainland: (2*s* +3) (*s*+d+4) cells. The island is initially a single cell at *s*+1 cells from the left, top, and bottom borders of the grid, and the mainland is at the right grid margin with two columns of cells. (d) Island size over time (in Myr). (e) Flow chart illustrating the sequence of simulated processes. Note that ecological and evolutionary processes were performed every time step, but island dynamics took place at much greater intervals (see panel d and ‘*Study design*’).

### State variables and scales

The model was grid-based (Fig. 1c) with a cell size of 1 km^2^, which was large enough to sustain viable tree populations yet small enough to distinguish between short-distance (≪ 1 km) and long-distance dispersal (Cabral & Kreft, 2012). Each island cell was assigned to an elevational level with an associated mean temperature. The model agents were populations, which were stage-structured (seeds, juveniles, adults) and measured in number of individuals. Populations belonged to species, which were conceptualized as specific combinations of autecological attributes, such as environmental requirements, dispersal ability parameters, and demographic traits (hereafter: species properties). Body mass and local temperature determined demographic transitions, mutation rates, the space exploited by an individual, carrying capacity, and time for speciation (see Appendix S1). These metabolic constraints accounted for the increase of metabolic rates with temperatures and their decrease with body mass (Brown *et al.*, 2004). The application of metabolic theory accounts for metabolic trade-offs related to energy allocation (e.g. survival vs. growth). Demographic transitions were germination, sexual maturation, reproduction, and density-independent mortality.

Unoccupied cell area was used as the interaction currency (Kissling *et al.*, 2012), for which populations competed. A cell could hold one population per species, but as many populations, and thus species, as there was area available. Consequently, species coexistence in a cell and on the island, and thus meta-communities and species assemblages, directly emerged from local resource competition (Cabral & Kreft, 2012). The state variables comprised the spatial distribution of seed, juvenile, and adult abundances of each species and the unoccupied area. Each time step was one year and a complete simulation ran over millions of years (Fig. 1d and *‘Study design’*). Eco-evolutionary processes took place every time step, whereas environmental events happened at longer intervals (Fig. 1e).

### Initialization

The model was initialized by reading in the simulation grid (Fig. 1c) and run specifications: mean annual air temperature at sea level (298 K, or 25 °C), size of the species pool (1000 species), and intervals for randomly drawing the species properties from a uniform distribution (see Appendix S1 for the list of parameters, interval values, and the justification of the values).

For each species, we randomly assigned the following properties: maximum cell suitability, optimum temperature, temperature amplitude, optimum island side, island side amplitude (environmental requirements), life form, mean dispersal distance, dispersal kernel thinness, strength of Allee effects, stage-specific body masses, and phenological ordering related to other species. Environmental requirements determined the distribution and quality of suitable habitat for each species. Optimum island side and island side amplitude represented requirements other than temperature affecting species distributions on islands, such as sunlight exposure or windward/leeward differences in precipitation (Whittaker & Fernández-Palacios, 2007). Temporal variation of island side and local temperature combinations accounted for changes in environmental heterogeneity, with maximum heterogeneity occurring at maximum island size. Depending on these environmental requirements, each species received a habitat suitability matrix, ***H*** (see Appendix S1). Seeds could only germinate in suitable cells.

A dispersal kernel ***D*** was initialized for each species by generating a two-dimensional, grid-based Clark’s 2Dt kernel, with two parameters (mean dispersal distance and dispersal kernel shape parameter) that described short- and long-distance dispersal, respectively (Clark *et al.*, 1999). The explicit consideration of long-distance dispersal is an advantage of Clark’s 2Dt (Clark *et al.*, 1999; Nathan & Muller-Landau, 2000) because it allows the simulation of both within-island and mainland–island dispersal processes.

On the island, the abundance matrices for adults, ***N_a_***, juveniles, ***N_y_***, and seeds, ***N_s_***, of each species as well as the matrix with the area occupied by all individuals, ***A_t_***, were initialized empty.

### Dispersal from mainland

At each time step, ten random species from each mainland cell *f* dispersed to the island. The seed bank for each species at island cell *i* was incremented by Poisson (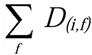 Uniform(1,10000)), where *D_(i,f)_* gives the dispersal probability per seed from cell *f* to cell *i*.

### Population update 1

Following the phenological ordering, we first calculated the area used by all individuals. Subsequently, abundances were sequentially updated by: i) turning juveniles to adults, ii) applying density-independent mortality to remaining juveniles, iii) germinating seeds, and iv) applying seed mortality (see Appendix S1 for equations). If germinating and maturing individuals in a cell surpassed the space available, excess individuals died. This accounted for density-dependent mortality through self-thinning, which is a difficult process to model (Reynolds & Ford, 2005; Wiegand *et al.*, 2008) but emerged in our simulations from space competition.

### Reproduction

The number of seeds produced by species *j* in cell *i* (*S_p(i,j)_*) was given by *N_a(i,j)_ R(N_a(i,j)_*), where *N_a(i,j)_* is the number of adults of species *j* in cell *i* and *R* a function describing per-capita density-dependent reproduction.

### Intra-island dispersal

Seeds of species *j* received in cell *z* and from source cell *i*, (*S_d(z,j)_*), were given by 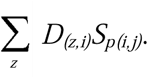.

### Mutation

We implemented a point-mutation process, which is a simple but efficient way to model cladogenesis (i.e. *in situ* speciation where two or more species evolve from a common ancestor, Rosindell & Phillimore, 2011). The number of mutated seeds was a Poisson random variate, whose probability was given by multiplying *S_d(z,j)_* with a metabolic mutation rate (Appendix S1). These genetically diverging individuals were initialized with random maximum cell suitability, optimum temperature, temperature amplitude, optimum island side, and island side amplitude (see *‘Initialization’*). The remaining species properties were randomly drawn within the ± 50% intervals around the values of the ancestral species to account for phylogenetic constraints. These evolving species properties allowed spatial and attribute divergence, which is commonly observed on islands.

### Speciation

We checked whether the time for genetically diverging individuals becoming a distinct species was reached (i.e. ‘protracted speciation’, Rosindell & Phillimore, 2011). We also included anagenesis (i.e. colonizers becoming distinct from the mainland form without *in situ* cladogenesis occurring). Both speciation modes depended on metabolic constraints to account for longer generations for larger body mass (Brown *et al.*, 2004). For anagenesis, the speciation time was counted from the island colonization event (no mutation required). However, every time mainland immigrants became adults, completion of anagenesis was delayed due to gene flow. The delay was arbitrary but varied metabolically and could re-start the speciation countdown (see Appendix S1). For cladogenesis, the speciation time was counted from the time step of the mutation event. Mutants surviving until their speciation time was completed were updated as cladogenetic endemics. The colonizer species giving origin to cladogenetic endemics remain either non-endemic under continuous gene flow or become anagenetic endemic.

### Population update 2

Abundance matrices were updated sequentially by applying density-independent mortality to adults and updating the seed bank after dispersal and mutation. These stochastic transitions followed respective metabolic transition rates (Appendix S1).

### Environmental dynamics

We considered the geo-climatic changes assumed on oceanic hotspot islands (Fig. 1b; Whittaker & Fernández-Palacios, 2007). To mimic volcanic island growth, each simulation started with a single cell that grew regularly by adding concentric belts of cells around the margins and by uplifting the interior belts. Thereafter, island size remained temporarily stable, followed by an erosion phase. At each erosion step, one belt of cells disappeared from the island margin, and the elevation of remaining cells decreased (Fig. 1b). Temperature decreased by 1 K for each elevational belt following uplift and increased by 1 K following erosion. We assumed an arbitrary length for each growth time step of 0.13 Myr (based on estimates for Madeira, see Appendix S1). We doubled this time length for the stable phase and erosion time steps to account for a slower erosion phase compared to the volcanic growth phase (Whittaker & Fernández-Palacios, 2007). Each simulation spanned 2.21 Myr. After every environmental event, ***H*** was recalculated for every species.

### Output

Output variables were ***N_a_, N_y_***, and ***N_s_***. Additionally, we recorded for every time step and cell species richness (total, non-endemics, and anagenetic as well as cladogenetic endemics), radiating lineages (species showing cladogenesis on the island) and species per radiating lineage. For the entire island, we counted colonization, speciation, and extinction events per time step.

### Study design

We designed two simulation experiments. First, we simulated the full model including all key processes. For each ecological level, we assessed multiple patterns and compared them to empirical data and theoretical predictions whenever possible. At maximum size, the simulated islands were 300 cells away from the mainland. The maximum island size was 11×11 cells and the mainland species pool comprised 1000 species. For this experiment, we simulated 20 replicate runs, each with a different mainland source pool.

The second experiment simulated exploratory scenarios in which we switched off processes to assess their relative role. Three replicate model runs were performed for each exploratory scenario. This replicate number allowed us to explore all scenarios, while still producing variability in the resulting time series. We simulated four scenarios: 1) without competition; 2) without metabolic constraints; 3) without environmental dynamics; 4) without speciation. Except for the scenario without metabolic constraints, replicate runs were based on the same source pool, thereby controlling for source pool-related variability. For the scenario without competition, we excluded competition with other source pool species by simulating each of 500 random species from the source pool in separate. However, competition was kept between evolving species. Otherwise, endemic richness would have increase exponentially. For the scenario without metabolic constraints, we switched off the body mass and local temperature control on demographic and evolutionary rates, which were then drawn independently (e.g. no trade-offs between reproduction and survival rates; Appendix S1). For the scenario without environmental dynamics, the island had a constant size of 7×7 cells. For the scenario without speciation, we switched off anagenesis, mutation, and cladogenesis. Given our focus on general trends and causal effects, we depicted variability of model simulations (e.g. 95% confidence envelopes) but we did not perform statistical tests when comparing scenarios because significance for minor differences emerges simply by increasing replicates (White *et al.*, 2014).

## RESULTS

The first simulation experiment generated temporally and spatially-explicit patterns spanning four different ecological levels. At the population level, the dynamics of population structure and species abundance at local scale and for the entire island emerged from the simulations (Fig. 2 a-f). Population structure (proportion of seeds, juveniles, and adults within the population) varied from mostly stable (Fig. 2d) to episodically changing due to island growth or erosion events (Fig. 2e-f) to ever changing (mostly due to declining adult abundances, Fig. 2c). Native non-endemics (from now simply non-endemics) tended to decrease in mean abundance (Fig. 2g), whereas anagenetic (Fig. 2h) and cladogenetic (Fig. 2i) endemics had more stable abundances over time.

**Figure 2.**
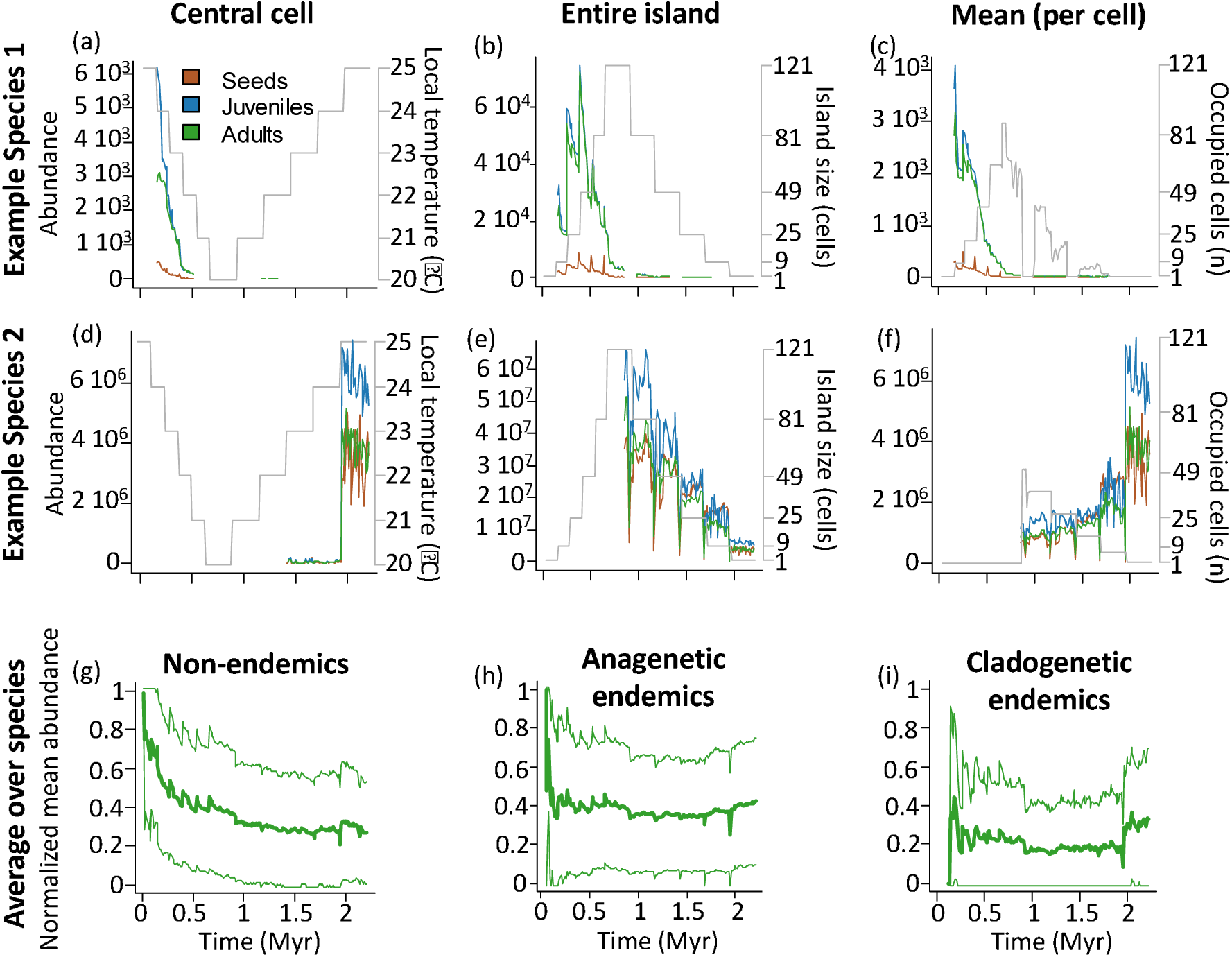
Population level temporal patterns. The top and intermediate rows show one exemplary species each, whereas the bottom row describes the abundance of adults normalized by dividing the time-series by the maximum abundance. (a-c) Seed, juvenile, and adult abundances (legend in a) in number of individuals of a non-endemic tree species adapted to lowlands and (d-f) of an endemic herb species adapted to lowlands (see Appendix S1 for detailed species properties). Panels (a) and (d) give local abundances in the central cell, (b) and (e) total island abundances, and (c) and (f) mean abundances (per occupied cell). The grey lines in (a-f) refer to the right y-axis: local temperature of the central cell (a, d), number of island grid cells (b, e), and occupied island grid cells (c, f). (g-i) Normalized mean cell abundances of (g) non-endemics, (h) anagenetic endemics (i) and cladogenetic endemics. Mean cell adult abundances were normalized by the highest value in the time-series per species. Thick lines in g-i indicated the average over replicates (n = 20) and species (n varies with replicate and time step), whereas thin lines indicate 95% CI (truncated at 0 and 1). The time-series were additionally averaged within time classes of 0.01 Myr.

At the species level, the spatial distribution of abundance and realized range changed over time and could strongly deviate from the distribution of potential habitat, with some species being able to survive only in sub-optimal environments (Fig. 3a). The species tended to have lower levels of range filling at intermediate to advanced island age, with cladogenetic endemics retaining higher range filling than anagenetic endemics and non-endemics (Fig. 3b).

**Figure 3.**
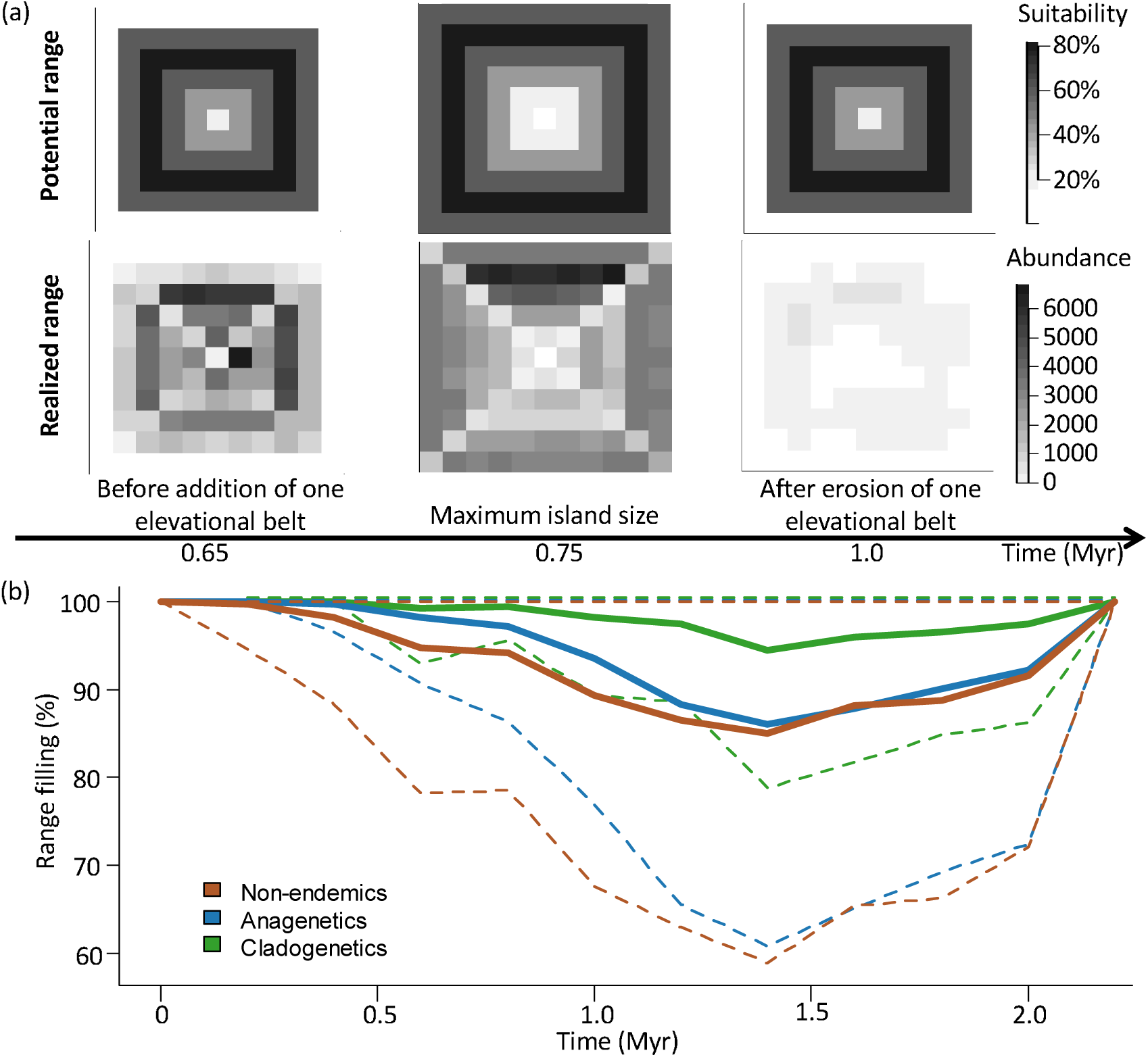
Temporal and spatial patterns at the species level. (a) Potential (top row) and realized (bottom row) range of a shrub species adapted to lowlands *(T_o_* = 24°C, *T_a_=* 4) at three different time steps: before (left column), during (middle column), and after (right column) the island has reached maximum size (see Appendix S1 for detailed species properties). (b) Range filling time series of range non-endemic species, anagenetic endemics, and cladogenetic endemics (n = 20; thick lines indicate means, thin dashed lines 95% CI). Time-series in (b) were averaged within each geological time step.

At the local community level, rank-abundance distributions followed a lognormal distribution (Fig. 4a). Local species richness, richness of cladogenetic endemics, and proportion of cladogenetic endemics increased over time (Fig. 4b-d, respectively). Species richness and cladogenetic endemic richness peaked at intermediate elevations (Fig. 4b-c, respectively), whereas the proportion of cladogenetic endemics was highest at low elevations (Fig. 4d).

**Figure 4.**
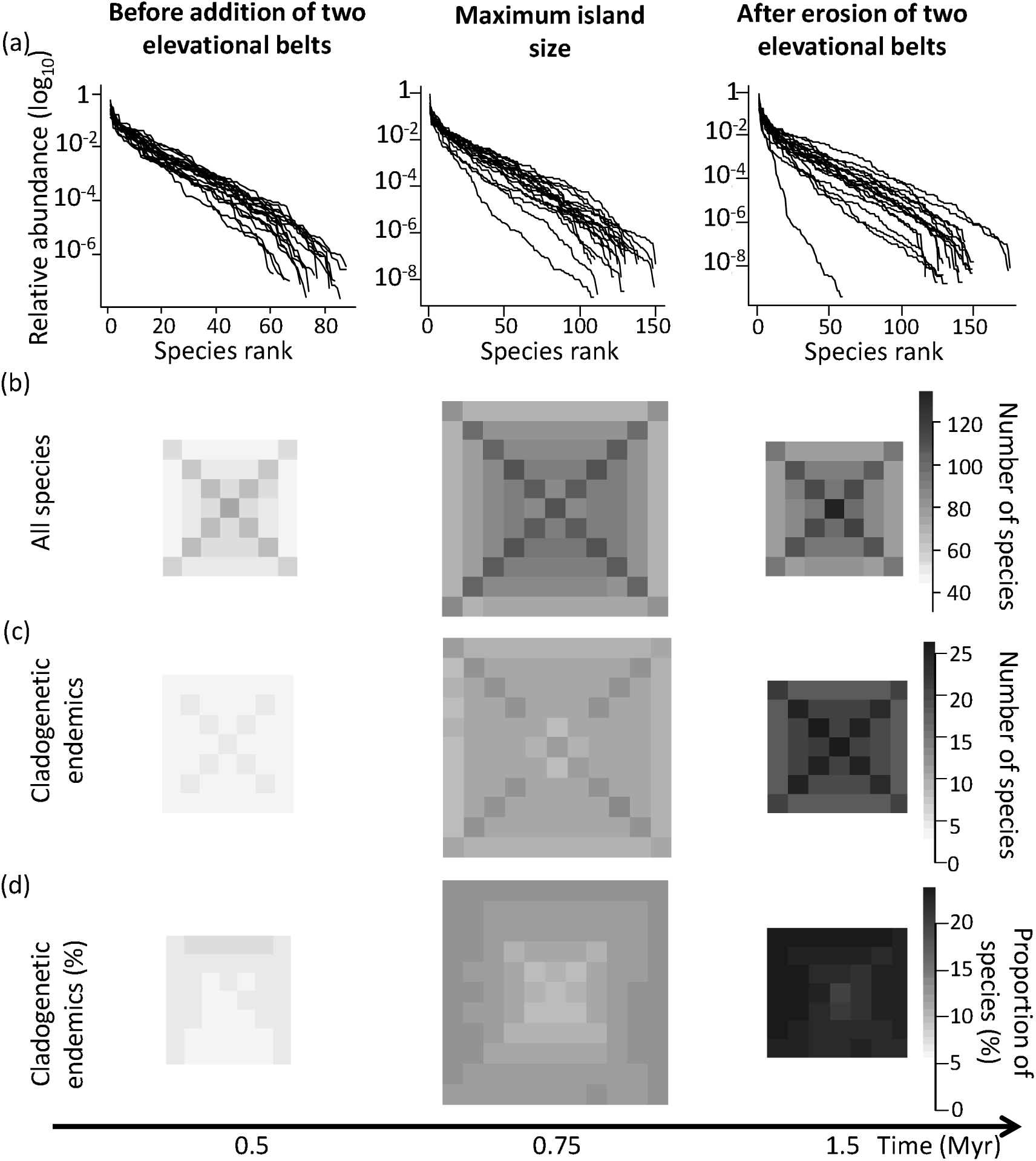
Temporal and spatial patterns at the community level. (a) Rank abundance plots of the central cell, (b) spatially-explicit species richness considering all species and (c) cladogenetic endemics, as well as (d) proportion of cladogenetic endemics at three different times steps: before (left panels), during (middle panels), and after (right panels) the island has reached maximum size. Rank abundances of each replicate are given by single lines in (a) and all of them fitted the lognormal distribution best when comparing AIC values for fitted logseries, lognormal and power law distributions to the abundance data (R package ‘sads’). Species richness and proportion of cladogenetic endemics were averaged over the 20 replicate runs.

At the level of the island-wide meta-community (from now on termed as assemblage), species–area relationships (SARs) were steeper during the growth phase than during the erosion phase (Fig. 5a-b). Species richness showed a humped trend and peaked at intermediate to advanced age (Fig. 5c), whereas the peaks for anagenetic and cladogenetic endemics lagged slightly behind (Fig. 5c). Colonization and extinction rates were humped and peaked at intermediate island ages (fig. 5d). This trend was also obtained for rates of anagenesis and cladogenesis, but it was two orders of magnitude lower than for immigration (Fig. 5e). Extinction of endemics was also humped, but with a very late peak (Fig. 5e). The number of radiating lineages, the number of species per radiating lineage, and trait richness all peaked at advanced island ages (Fig. 5f, g). However, species packing showed an initial steep decrease followed by a humped trend over time (Fig. 5h). For the cladogenetic endemics, the mean pairwise trait distance to their ancestral species showed a very shallow humped trend over time, whereas the distance to all species increased almost for the entire island lifespan, decreasing only at very advanced island age (Fig 5i).

**Figure 5.**
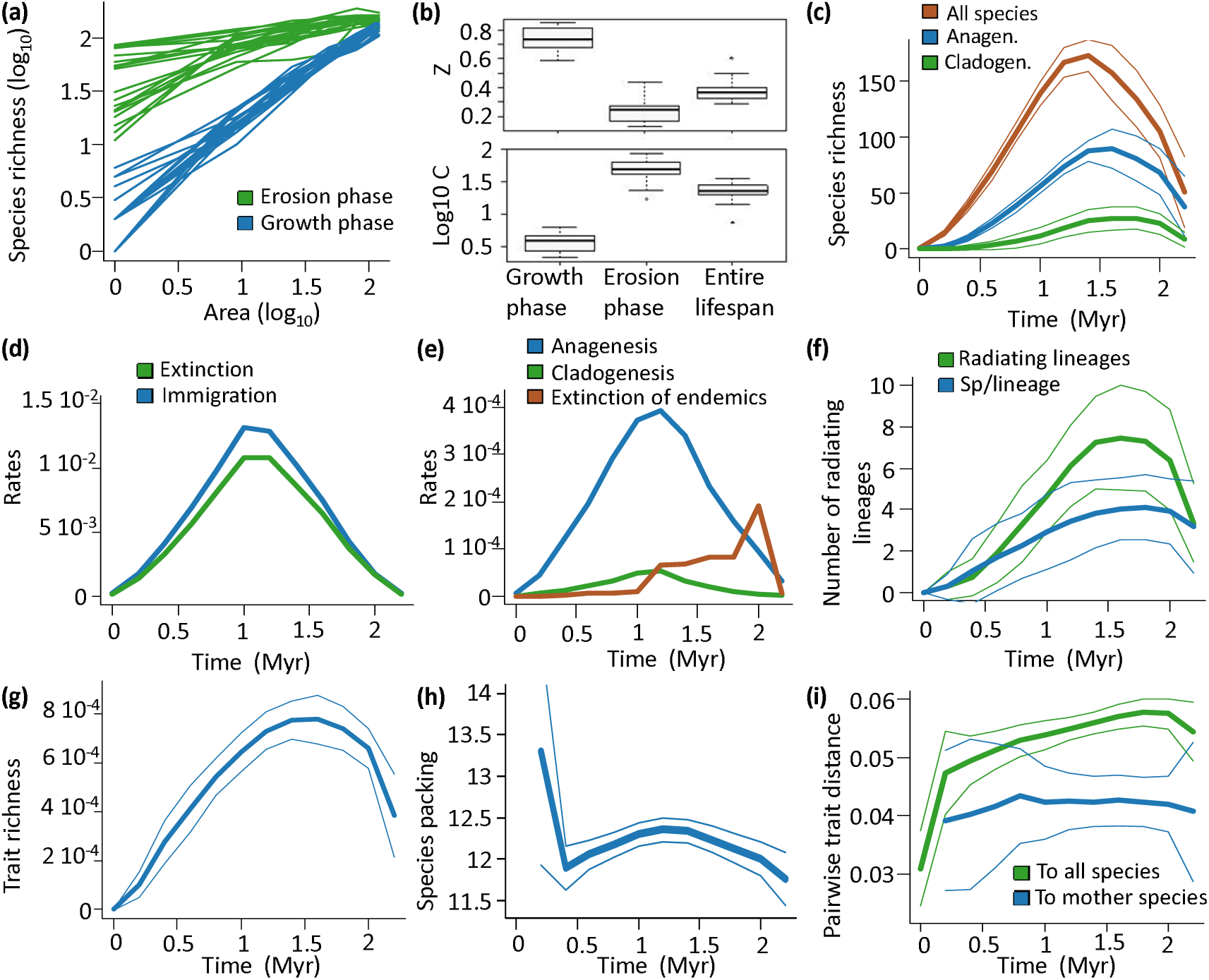
Temporal patterns at the assemblage level. (a) Log-log species-area relationships (SARs) for growth and erosion phases (lines represent replicates), (b) slope *z* and intercept logC of estimated power-law SARs for growth and erosion phases as well as the entire island geological lifespan, (c) species richness over time, (d) colonization and extinction rates over time, (e) anagenesis, cladogenesis, and extinction rates of endemics over time, (f) number of radiating lineages and species per radiating lineage over time, (g) trait richness (volume of the convex hull of the multivariate space considering all species properties) over time, (h) species packing (number of species per trait richness unit) over time, (i) pairwise trait distance of cladogenetic endemics to their mother species and to all other species over time, averaged per trait and per species pair. Thick lines in (c-i) indicate average within each environmental time step (see *‘environmental dynamics’* in the methods) and over 20 replicates, with 95% CI (thin lines, omitted in panels d and e for visual clarity).

The second simulation experiment, namely switching off main processes, revealed patterns diverging from the first, full model experiment (Fig. 6). At the population level, population structure was generally stable in the scenario without competition (Fig. 6a vs. Fig. 2b), whereas population structure was highly oscillatory without metabolic constraints (Fig. 6b). Without environmental dynamics, species colonized earlier, but were subjected to extinction as in the full model (Fig. 6c). Without speciation, colonizers decreased in abundances over time, but survived the entire simulation (Fig. 6d). Without competition, range filling was lower at intermediate island age and for cladogenetic species (Fig. 6e vs. Fig. 3b), whereas without metabolic constraints, range filling was variable, with non-endemics going extinct at advanced island age (Fig. 6f). Without environmental dynamics, range filling was highest for cladogenetic endemics, but showed stable dynamics after the initial colonization period (Fig. 6g). Without speciation, species maintained high levels of range filling (Fig. 6h). In the scenario without competition, the proportion of cladogenetic endemics was extremely high at advanced island age, particularly in the lowlands (Fig. 6i vs. Fig. 4d right panel). In the scenario without metabolic constraints, the proportion of cladogenetic endemics was also high, but without spatial structure (Fig. 6j). Moreover, without environmental dynamics, the proportion of cladogenetic species peaked at low and high elevations (Fig. 6k), whereas endemics were understandably absent without speciation (Fig. 6l). At the assemblage level, species richness showed a humped trend over time and was dominated by endemics, with very high values without competition (Fig. 6m vs. Fig. 5c) and very low values without metabolic constraints (Fig. 6n). Without environmental dynamics, total species richness on the island tended towards equilibrium but the number of cladogenetic species continued to increase (Fig. 6o), whereas without speciation, species richness showed a humped trend similar to, but with lower values and less variation than, the full model (Fig. 6p vs. Fig. 5c).

**Figure 6.**
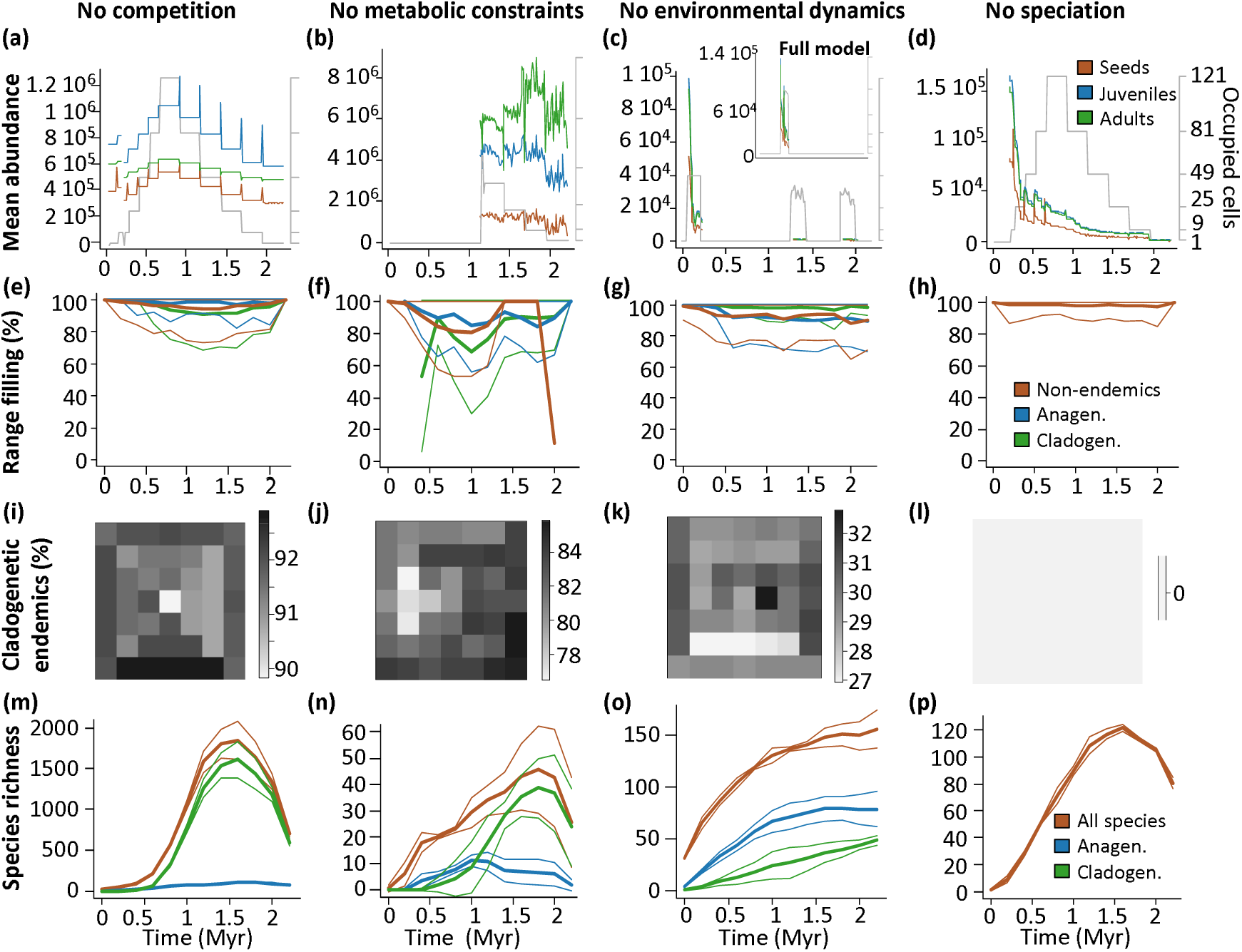
Evaluation of the model structure across patterns at different ecological levels (rows) by switching off key processes (columns). (a-d) Population dynamics of an example species, given by mean abundances (per occupied cell). (e-h) Overall range filling dynamics. (i-l) Proportion of cladogenetic endemics at advanced island age, 1.5 Myr. (m-p) Total species richness dynamics of exploratory scenarios with no competition (left column), no metabolic constraints (middle left column), no environmental dynamics (middle right column) and no speciation (right column). Colour legends differ between rows: legends in (c) for population dynamics, in (h) for range filling dynamics, and in (p) for total richness dynamics. The panels (a), (c), and (d) illustrate the population dynamics of one example non-endemic shrub species adapted to intermediate elevations that survived in these three scenarios and in the full model (inset in c). Panel (b) illustrates the population dynamics of one example cladogenetic endemic herb species adapted to lowlands (see Appendix S1 for detailed species properties). Grey lines and right y-axis in (a-d) indicate the number of occupied cells. The other panels illustrate values averaged over replicates (n = 3) and within each geological time step (95% CI given as thin lines for range filling and total species richness dynamics).

## DISCUSSION

### Population level

In the full model, population structure varied according to local environment, species evolutionary origin (non-endemic, anagenetic, or cladogenetic), and island age (Fig. 2). The high relative abundances, particularly of non-endemics (Fig. 2g), on young islands can be explained by lower competition due to low species number. Population establishment decreases with island age, a pattern which has been observed for biological invasions on islands worldwide (Kueffer *et al.*, 2010). Interestingly, endemics were better able to sustain stable or increasing populations compared to non-endemics (Fig. 2g-i). Empirical data comparing abundances of endemic and non-endemic plant species are rare, but evidence from pollination networks on oceanic islands suggests that endemics tend to be generalists and to have higher relative abundances than non-endemics (Olesen *et al.*, 2002). Moreover, endemic arthropods in the Azores also tended to show higher densities and occupancy than native non-endemic species (Gaston *et al.*, 2006). Strikingly, without competition, species could sustain high and stable abundances (Fig. 6a), whereas without speciation, populations decreased over time, but could survive longer than in the full model (Fig. 6d). These results confirm our first hypothesis that competition regulates populations, particularly considering the colonization and evolution of competitors.

### Species level

Under the full model experiment, the sharp decrease in normalized abundances after the initial island growth phase (Fig. 2g - i) translated into a general decrease in range filling at the species level at intermediate to advanced island ages (Fig. 3). Even if species could fill their entire potential range, the realized abundance distribution could diverge from the distribution of habitat suitability, particularly at intermediate island age (Fig. 3a). Such divergence can be explained by interspecific competition, which decreases abundances and may shift abundance peaks to sub-optimal environmental conditions, as reported for plants and animals (McGill, 2012; Wisz *et al.*, 2013). With decreasing island size, extinction of poorer competitors increased range filling of surviving species (Fig. 3b). Remarkably, cladogenetic endemics were better able to fill their potential range compared to non-endemics and anagenetic endemics, which indicates high selective pressure on species to cope with competition. Because anagenetic endemics did not change their niche upon speciation, their range filling dynamics were coherently comparable to non-endemics.

The scenario without competition revealed similar trends, but with higher range filling compared to the full model, with cladogenetic endemics showing the lowest values (compare Fig. 6e with Fig. 3b). The lowest values for cladogenetic endemics reflected the persistence of competition between endemics in this scenario (see Materials and Methods) and that the number of endemics was very high (Fig. 6m). Hence, the lower range filling of cladogenetic endemics can be explained by an almost unconstrained diversification. Accordingly, in the scenario without speciation, and thus without competition with endemics, non-endemics achieved high range filling throughout the simulation (Fig. 6h). In contrast, without metabolic constraits, super-dominant species evolved, leading to low range filling followed by extinction of non-endemics (Fig. 6f). Together, these results confirmed our first hypothesis that competition is necessary to regulate range dynamics (see also Singer *et al.*, 2016; Urban *et al.*, 2016), mostly via competition between non-endemic and endemic species.

### Community level

The full model produced realistic rank abundance distributions with only few dominant species (Fig. 4a). The general lognormal shape of the rank abundance plots were consistent with a recent meta-analysis (Ulrich *et al.*, 2010). Moreover, species abundance distributions could be described by the recently introduced gambin distribution (Appendix S2; Matthews *et al.*, 2014). When assessing the spatial distribution of local species richness, the mid-elevation peaks (Fig. 4b-c) reflected the random distribution of temperature niches of the source pool, with most ranges overlapping in mid-elevations due to mid-domain effects (Colwell & Lees, 2000; Cabral & Kreft, 2012). Accordingly, cells between island sides had higher richness reflecting ecotones caused by overlapping of ranges of island side specialists. Remarkably, the percentage of cladogenetic endemics was highest at lower elevations (Fig. 4d). On real islands, species richness often peaks at low or mid-elevations (Sanders & Rahbek, 2012; Seipel *et al.*, 2012). The percentage of single-island endemics, in contrast, tends to peak at high elevations (Steinbauer *et al.*, 2012), which might be due to higher isolation, lower competition, and lower gene flow in high-elevation environments (Steinbauer *et al.*, 2012). In our simulations, intermediate elevations were the least isolated environments due to the random assignment of temperature niches, which resulted in more species overlapping their ranges at these elevations. Therefore, the higher proportion of cladogenetic endemics in the lowlands in the full model supports the isolation effects because these elevations were suitable for a lower number of species compared to intermediate elevations. The low endemism at high elevations can then be explained by time and area effects, considering that high elevations had the smallest area and had less time to accumulate species. In fact, the scenario without environmental dynamics showed a high proportion of cladogenetic endemics at both low and high elevations (Fig. 6k).

Beyond area, time, and isolation, environmental variables also affected endemism via increasing mutation and speciation rates with temperature (Allen *et al.*, 2006). Very high values and no evident spatial structure of cladogenetic endemism in the scenario without metabolic constraints (Fig. 6j) confirmed that these constraints influence the spatial structure of speciation and local communities, supporting our second hypothesis. These constraints prevent super-dominant species (e.g. species with high reproductive and survival rates combined with low resource requirements □ Brown *et al.*, 2004). Moreover, metabolic constraints interact with competition to regulate optimal environments for speciation, indicated by the mid-elevation peaks in endemic richness in the full model. This happens because higher speciation rates in the lowlands can be prevented by higher resource requirements (Brown *et al.*, 2004) and thus stronger interspecific competition, despite higher mutation rates (Allen *et al.*, 2006) as well as higher resource availability (i.e. larger area). Consequently, mid-elevations represent the best balance between higher mutation rates and resource availability in the lowlands vs. lower competition pressure in the highlands.

### Assemblage level

Species richness of entire islands closely tracked environmental dynamics (Fig. 5a-b). Slope values of power-law SARs considering the entire island life-span matched remarkably well with real-world SARs of oceanic islands (mean *z* = 0.39 in Fig. 5b compared to *z* = 0.38 in Triantis *et al.*, 2012). Moreover, the SAR intercepts reflecting the average species density (mean *c* = 1.4, Fig. 5b) were also comparable to intercepts reported for plants (*c* = 1.6 in Triantis *et al.*, 2012), but larger than intercepts reported for oceanic islands (*c* = 0.6; Triantis *et al.*, 2012). Such low reported intercepts are, however, closer to the intercepts obtained when considering only the growth phase (*c* = 0.55; Fig. 5b), when the islands start devoid of life. In real world, low intercepts (i.e. low richness per unit area) are found for area-demanding taxa (e.g. vertebrates and trees) and on very small, low-lying islands, such as atolls. These islands are also subject to frequent disturbances (Morrison, 2010), which are not simulated. Nevertheless, the obtained differences in SARs indicate that future studies should account for the geological phase of the islands.

Species richness and endemic richness (Fig. 5c) followed the humped trend over time predicted by the GDM (Whittaker *et al.*, 2008; Hortal *et al.*, 2010; Borregaard *et al.*, 2016). This hump was absent in the scenario without environmental dynamics (Fig. 6o), confirming our third hypothesis that the environmental processes drive the temporal dynamics of species richness. Without environment processes, the seemingly stable species richness (ca. 160 species, Fig. 6o) is in line with the ETIB assumption of static islands (MacArthur & Wilson, 1963). However, whereas the species richness stabilizes through a dynamic equilibrium between colonization and extinction in ETIB, our results represent a dynamic balance between colonization, speciation, and extinction with a almost constant disequilibrium caused by environental dynamics. Remarkably, without ecological and evolutionary processes the humped trend was retained but richness values were affected (Fig. 6m-n, p). In particular, species richness reached high values without competition (Fig. 6m), whereas it was very low without metabolic constraints (Fig. 6n). In both scenarios, island floras were dominated by endemics, suggesting that competition and metabolic constraints affect evolutionary processes and species assemblage composition, although their impacts are assumed to mostly affect richness locally (but see Waters *et al.*, 2013; Pedersen *et al.*, 2014).

The humped distribution of species richness over time in the scenario without speciation rejected our forth hypothesis that speciation is necessary to generate this pattern. Our results indicate that the humped richness can emerge through colonization and extinction rates alone and without involving speciation rate. In the full model, colonization and cladogenesis rates indeed followed the theoretical predictions of decreasing colonization and humped speciation rates for a hotspot geological trajectory (Fig. 5d-e; Whittaker *et al.*, 2008; Borregaard *et al.*, 2016). Moreover, within cladogenesis, the numbers of radiating lineages and species per radiating lineage over time (Fig. 5f) were also consistent with theoretical predictions (Whittaker *et al.*, 2008; see also Cabral *et al.*, submitted). This indicates that diversification continues after volcanic activity has ceased and area has decreased (Whittaker & Fernández-Palacios, 2007). Furthermore, the humped extinction (Fig. 5d) and anagenesis (Fig. 5e) rates were consistent with GDM predictions (Borregaard *et al.*, 2016). Overall extinction rates followed the expected humped colonization trends (Fig. 5e; Borregaard *et al.*, 2016) and reached a dynamic equilibrium only on very old islands akin to ETIB predictions (similar extinction and immigration rates in Fig. 5d; MacArthur & Wilson, 1963). Interestingly, the much delayed peak extinction of all compared to that only of endemic species (compare Figs 5d-e) indicates that endemic species might be less susceptible to extinction (i.e. better adapted to in situ conditions) than non-endemic species during early stages of island erosion.

Trait richness followed the humped trend of species richness, consistent with the positive relationship between trait richness and species richness (Fig. 5c,g; Petchey & Gaston, 2002; Carnicer *et al.*, 2012). We found initially high values in species packing followed by a sharp decrease (Fig. 5h), possibly reflecting strong environmental filtering, indicating that only species with similar trait syndromes (e.g. lowland-adapted, good dispersing herbs) might be able to colonize young islands. As islands gain in environmental heterogeneity, new colonizers and evolving endemics increased the trait space, causing the sharp decrease in species packing (Fig. 5h). Thereafter, species start to fill the trait space, closely tracking island area. However, species packing was not random, as cladogenetic endemics were selected to fill the environmental space away from the co-occurring species thereby avoiding niche overlap and competitive exclusion (e.g. trait dispersion or character displacement; Mizera & Meszéna, 2003), while their trait distance to the ancestor was phylogenetically constrained (Fig. 5i). This indicates complex interactions between trait, demographic, and diversification dynamics (see Carnicer *et al.*, 2012 for a review) and a strong influence of competition on trait selection and evolution.

### Model properties, limitations and potentials

The ability of the model to simultaneously generate multiple patterns across different ecological levels provides opportunities for cross-scale validation (Grimm & Railsback, 2012). Other process-based island models (Kadmon & Allouche, 2007; Hortal *et al.*, 2009; Rosindell & Phillimore, 2011; Rosindell & Harmon, 2013; Valente *et al.*, 2014, 2015; Borregaard *et al.*, 2016) have fewer parameters and thus lower complexity. However, these models tend to simulate biogeographical processes directly (e.g. colonization, extinction, and speciation) and are spatially-implicit. Our approach, in contrast, describes the same biogeographical patterns emerging from population-level processes in a spatially-explicit context. The unrealistic patterns we obtained when switching off core processes support the importance of these processes and justify model complexity. Additionally, the explicit representation of space and environmental heterogeneity of our approach facilitates a niche-based framework, which is fundamental for testing island biogeography theory given the role of habitat heterogeneity and niche opportunities for speciation (Whittaker *et al.*, 2008).

Limited availability of empirical data generally hampers model validation, parameterization, and quantification of model uncertainty (Jeltsch *et al.*, 2008; Dormann *et al.*, 2012). The hierarchical structure of our model allows the use of different data types of to calibrate the model and to evaluate different emergent patterns (Wiegand *et al.*, 2003). For example, estimates of demographic rates can be used to fit metabolic functions (Schurr *et al.*, 2012) and abundance distributions to fit demographic functions (Cabral & Schurr, 2010). Simulating large and species-rich islands might be computationally unfeasible, but within feasible computational scenarios data scarcity can be overcome with pattern-oriented modelling by using emergent patterns to calibrate unknown parameters and preventing error propagation (Wiegand *et al.*, 2003; Grimm & Railsback, 2012).

We used theoretical predictions and empirical data for evaluation of model structure and validation of the full model. A range of emergent patterns followed well-documented empirical trends and relationships. Namely, these were rank-abundance distributions (Ulrich *et al.*, 2010), relationship between proportions of endemic species and environmental isolation (Steinbauer *et al.*, 2012), species–area relationships (Triantis *et al.*, 2012), species richness and endemism over time (Whittaker *et al.*, 2008; Cameron *et al.*, 2013). Such cross-scale and cross-ecological level validation suggests that our model produces generalizable predictions over a wide range of systems (see Evans *et al.*, 2013 for a review on generality of complex models). Some of the patterns generated currently lack empirical data for evaluation and thus constitute predictions to be tested in future studies (e.g. humped trait diversity over time, Fig. 5g). Additionally, in all predictions (with and without available comparable empirical data), our model inherently integrates variability by considering demographic, colonization, extinction and speciation stochasticity via multiple replicate runs. Finally, parameter and model uncertainties can be addressed by varying scenarios (e.g. different isolation scenarios, Cabral *et al.*, submitted) and model structure (Fig. 6). Therefore, data limitation should not prevent the exploration of relevant processes in simulation models (Evans *et al.*, 2013).

### Model implications and conclusions

Our modelling results show that understanding biodiversity dynamics requires the consideration of many different ecological, evolutionary, and environmental processes. In an island biogeography-related context, the novelty of our approach is that it simulates processes at the level of the individuals and populations in a stochastic, niche-, and metabolism-based framework. This framework leads to biogeographical dynamics emerging at large spatiotemporal scales. Our approach thus unifies mechanistically multiple ecological and evolutionary theories with island biogeography theory. Besides confirming several predictions of island biogeography theory, the integration of eco-evolutionary and environmental processes pinpointed interesting divergences and provided insights into less studied patterns and process interactions. We therefore argue that process-based models hold a great potential to serve as ‘virtual, long-term field stations’ in biogeography.

## ACKNOWLEDGEMENTS

J.S.C. acknowledges funding from the German Research Foundation (DFG; SA 2133/1-1) and from sDiv, the Synthesis Centre of iDiv (DFG FZT 118). H.K. was supported by the DFG through the German Excellence Initiative. K.W. was partly funded by the State of Lower Saxony (Ministry of Science and Culture; Cluster of Excellence “Functional Biodiversity Research”). We thank Albert Phillimore, James Rosindell, Kostas Triantis, Robert Whittaker, Yael Kisel, Gunnar Petter, Anke Stein, Patrick Weigelt, Carsten Meyer for feedback on earlier manuscript versions and a special thanks to Volker Grimm for the encouragement and feedback on the project.

## SUPPORTING INFORMATION

Additional supporting information may be found in the online version of this article:

**Appendix S1** Detailed model description.

**Appendix S2** Gambin fit to species abundance distributions.

## BIOSKETCH

**Juliano Sarmento Cabral** is interested in processes and factors influencing species and biodiversity dynamics across spatio-temporal scales. His research includes processes determining spatial and temporal distribution of tropical epiphytes, species ranges, island plant diversity as well as global species richness and endemism patterns.

**Kerstin Wiegand** is interested in the role of space for population dynamics, interspecific interactions, and biodiversity. The methodological emphasis of her research is on (spatially explicit, agent-based) simulation models and spatial statistics.

**Holger Kreft** has a broad interest in biogeographical and macroecological patterns, particularly gradients in species richness and endemism. His research includes analyses of plant and vertebrate diversity, island and conservation biogeography.

## Author contributions

J.S.C. and H.K. designed the study, with input from K.W.; J.S.C. implemented and simulated the model; J.S.C. led the analyses and writing, with input from all co-authors.

